# A versatile compressed sensing scheme for faster and less phototoxic 3D fluorescence microscopy

**DOI:** 10.1101/125815

**Authors:** Maxime Woringer, Xavier Darzacq, Christophe Zimmer, Mustafa Mir

## Abstract

Three-dimensional fluorescence microscopy based on Nyquist sampling of focal planes faces harsh trade-offs between acquisition time, light exposure, and signal-to-noise. We propose a 3D compressed sensing approach that uses temporal modulation of the excitation intensity during axial stage sweeping and can be adapted to fluorescence microscopes without hardware modification. We describe implementations on a lattice light sheet microscope and an epifluorescence microscope, and show that images of beads and biological samples can be reconstructed with a 5-10 fold reduction of light exposure and acquisition time. Our scheme opens a new door towards faster and less damaging 3D fluorescence microscopy.

**OCIS codes:** (110.1758) Computational imaging; (170.2520) Fluorescence microscopy; (170.6900) Three-dimensional microscopy.

## 1. Introduction

Imaging fluorescently labeled biological structures with high spatio-temporal resolution requires judicious compromises between the conflicting goals of achieving high signal-to-noise ratio (SNR) and temporal resolution while keeping the excitation power low to minimize photobleaching and phototoxicity. For example, to obtain a higher SNR, one can either increase the exposure time, thereby reducing imaging speed, or increase the illumination power, thereby increasing photodamage. These tradeoffs are further exacerbated in 3D imaging, which is often required in biological applications, such as calcium imaging in neurons or transient mitotic events in a developing embryo. Traditionally, 3D microscopy images are obtained by sequentially acquiring 2D images of individual focal planes, where axial spacing is dictated by the Nyquist sampling criterion to achieve optimal spatial resolution in all dimensions. As a consequence, hundreds or thousands of planes are needed to image samples 10-1,000 *μm* thick, dramatically increasing acquisition time and light exposure. Although light sheet illumination considerably reduces photodamage and allows prolonged imaging of living cells in 3D, the required hardware systems are often costly and scarce, and the acquisition time for each volume still remains constrained by the Nyquist criterion [1–3].

The field of compressed sensing, introduced over a decade ago, offers an avenue to overcome these limitations [4–7]. Compressed sensing leverages the fact that natural images are highly non-random and harbour intrinsic redundancies, which can be formulated as sparsity in an appropriate linear basis [8]. This sparsity can be exploited in order to reconstruct images from fewer measurements than specified by the Nyquist criterion, provided that measurements are taken in an appropriate manner. Compressed sensing has been successfully applied in diverse imaging applications in fields including astronomy [9] and magnetic resonance imaging [10], where it has enabled a considerable increase in acquisition speeds.

In biological microscopy, compressed sensing should in principle enable similar benefits in reducing acquisition time and light exposure without compromising SNR [11]. However, despite several proof of concepts, fluorescence microscopy has benefited relatively little from compressed sensing approaches in practice. One reason for this is that most compressed sensing strategies proposed to date require considerable modifications of the optical system, an important impediment for application on routinely used microscopes [12–17]. We note however, that a compressed sensing scheme without modification of the light path was recently used in confocal laser scanning microscopy to achieve a 10-15 fold speedup in 2D imaging [18].

Here we introduce a compressed sensing scheme for 3D fluorescence imaging that relies on compression along the optical axis (*z* axis) and is applicable to a large range of fluorescence modalities without modification of the optical path. We show that for a given SNR, our method can reconstruct a *z* stack from a 2-10 times faster acquisition than traditional plane-by plane imaging with Nyquist sampling. For dynamic microscopy of live samples, this approach opens the door to either lower excitation power and photodamage (at constant acquisition speed and SNR) or to higher temporal resolution (at constant excitation power and SNR).

In Section 2, we first describe our method conceptually, starting with a brief reminder of the basics of compressed sensing. In Section 3, we present results on simulations. Implementation on a lattice light sheet microscope and a conventional epifluorescence microscope are demonstrated in Sections 4 and 5 respectively. Sections 6 and 7 provide a brief discussion and conclusion.

## 2. Method

### 2.1. Basics of compressed sensing

Compressed sensing is based on the realization that under certain (broad) conditions, natural signals such as images can be reconstructed from a smaller number of measurements than prescribed by Nyquist sampling. If *X* is a (vectorized) image of size *N*×1 and **A** a known *M* × *N* matrix that transforms *X* into a signal *Y* = **A***X* of smaller size *M* × 1 (*M* < *N*), then the goal is to recover *X* (or a good approximation thereof) from *Y*. The matrix **A**, which is independent of the data, specifies how the *N* pixels of the image are scrambled into the *M* “compressed” measurements and is called the sensing (or measurement) matrix.

In order to recover *X* from *Y*, compressed sensing reconstruction algorithms exploit the structural redundancy of images. In the simplest setting, it is assumed that the image *X* is sparse, i.e. that the number of non-zero values, *K* = ||*X*||_*ℓ*0_ is small (*K* << *N*), or that *X* can be represented sparsely in a suitable basis, i.e. that *X* = Ψ*α*, where Ψ is an invertible (e.g. orthonormal) *N* × *N* matrix and α is a sparse vector of size *N* × 1. Note that this setting can easily be adapted to incorporate a redundant dictionary Ψ of size *W* × *N* with *W* > *N* instead of an orthogonal basis, allowing for improved reconstructions [32, 33]. The reconstruction algorithms aim to determine the sparsest representation *α* consistent with the data, i.e. such that *X* = **A**Ψ*α*. While minimizing the *ℓ*_0_ norm to enforce sparsity is NP-hard and computationally unfeasible, minimizing the *ℓ*_1_ norm 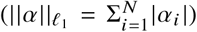 leads to computationally tractable optimization algorithms that recover the exact solution.

In practice, images are noisy and only approximately sparse, therefore compressed sensing algorithms seek to recover approximations of *X* by determining the sparsest representation *α* such that *Y* ≈ **A**Ψ*α*. For additive gaussian noise, this is typically done by solving the optimization problem: *α*\* = arg min_*α*_ *F*(*α*) for objective functions *F*(*α*) such as:

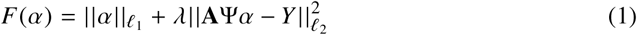

where *ℓ*_2_ is the Euclidian norm and *λ* a Lagrange multiplier.

Under suitable conditions for **A**, such as the restricted isometry property (which is fulfilled in particular for random Gaussian matrices), it was shown that a good approximation of the *N* values in *α* can be recovered from a number of compressed measurements *M* = 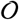 (*K* log(*N/K*)), which can be much smaller than *N* [19–22]. The reconstructed image is then simply obtained as *X*\* = Ψ*α*\*. For images corrupted by Poisson noise, the appropriate objective function becomes:

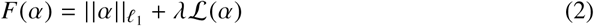

and its minimization is subject to the positivity constraint: Ψ*α* ≥ 0, where 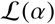 designates the negative Poisson log-likelihood 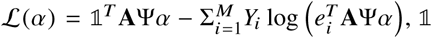 is a *M* × 1 vector of ones, *e_i_* is the canonical basis vector *i* and the *T* superscript denotes transposition [24]. A variety of efficient algorithms for compressed sensing recovery have been proposed, mostly for Gaussian noise, but also for Poisson noise [23, 24]. See [7, 8, 25] for in-depth introductions to sparsity and compressed sensing.

### 2.2. Axially compressed imaging scheme

The traditional way to image a 3D volume is to successively scan the focal plane of the microscope along the *z* axis in a step-wise fashion, with spacing Δ*z*, and acquire a 2D image at each *z* = *k*Δ*z* position (*k* = 1 … *N*). We hereafter refer to this imaging scheme as *plane-by-plane* acquisition (Figure 1.a). In this scheme, the focal plane position is given by:

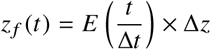

where Δ*t* is the camera exposure time and *E*(*x*) is the next smallest integer to *x*. The spacing Δ*z* is usually dictated by the point spread function (PSF) width along the *z*-axis (Nyquist sampling). In plane-by-plane imaging, the *k*-th camera frame, 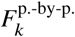, *k* = 1 … *N* carries information from the *z* = *k*Δ*z*-plane only and is given by:

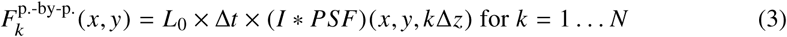

where *I* (*x, y, z*) designates the 3D distribution of fluorophores in the sample, *L*_0_ is the laser intensity, *PSF* is the 3D PSF of the microscope and * stands for convolution. In this setting, the acquisition time for a full 3D *z*-stack with *N* focal planes is *N*Δ*t* and the light dose received by the sample is *NL*_0_Δ*t*.

**Fig. 1.**
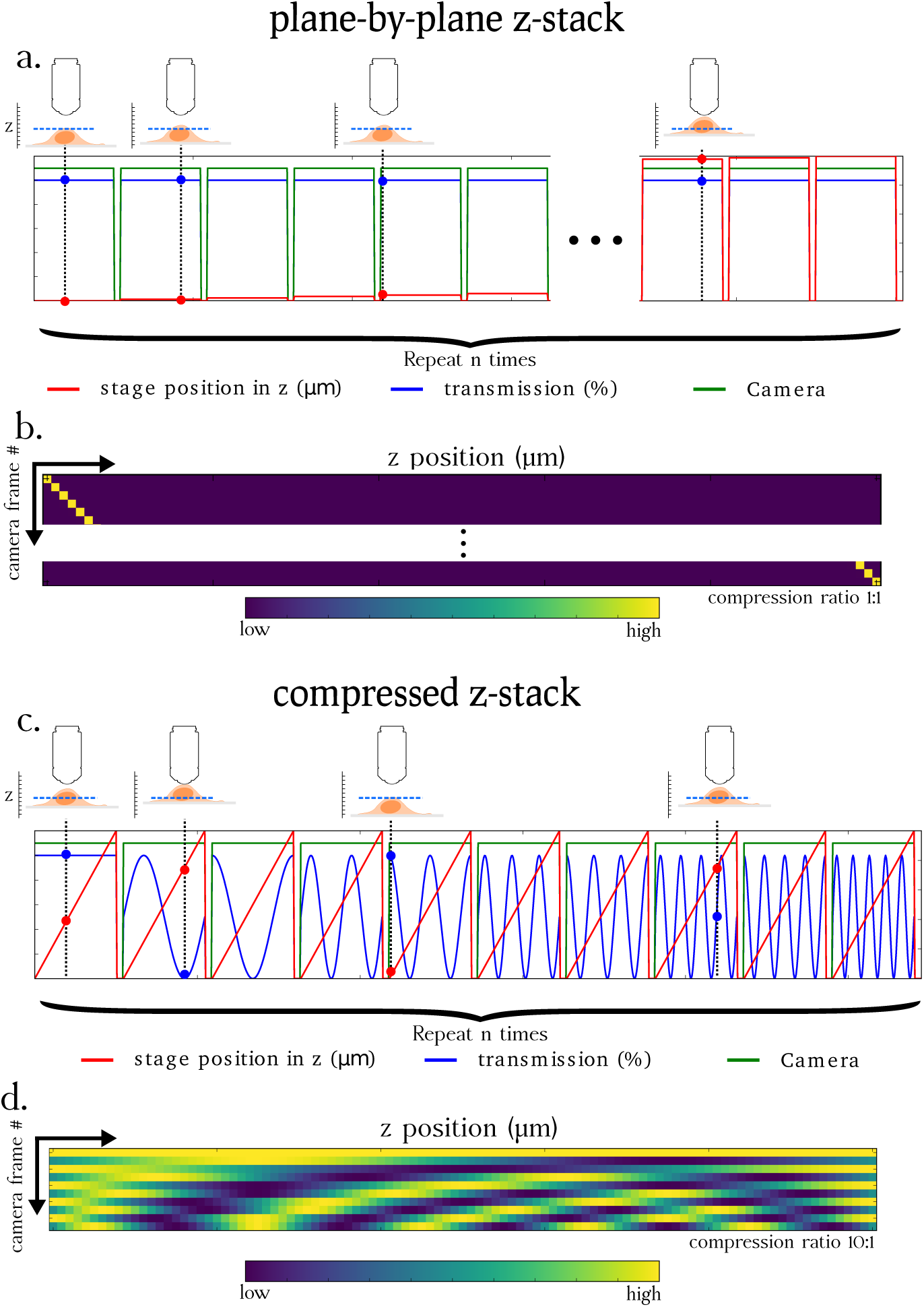
Principle of the proposed 3D compressed imaging method compared to traditional 3D plane-by-plane imaging. **(a)** Plane-by-plane imaging: for each camera frame (green curve indicates if the shutter is open or closed), one plane of the sample (red curve indicates *z* position) is illuminated at a constant laser intensity (blue curve shows transmission percentage of the excitation light source). The process is repeated for each plane (*N* = 101 times) to acquire a *z*-stack. The illustrations above the chart exemplify the position of the stage and the illumination intensity at various time points. This imaging scheme can be represented as the application of a square diagonal measurement matrix **A** as shown in **(b)**: for each camera frame (row), only one *z* plane is illuminated (column). **(c)** Axially compressed imaging: the stage continually sweeps through the entire axial range while the illumination is modulated to create a specific axial light pattern. In this scheme, multiple planes of the sample are illuminated during a single camera exposure frame. This process is repeated *M* = 10 < *N* times with different light patterns, thus performing an optomechanical implementation of a compressed measurement matrix, as shown in **(d)**.

In the compressed sensing imaging scheme proposed here, the axial dimension of a 3D stack is acquired in a compressed fashion, such that the *k*-th acquired frame no longer contains information from the *z* = *k*Δ*z* position only, but is a linear combination of information from multiple z-planes:

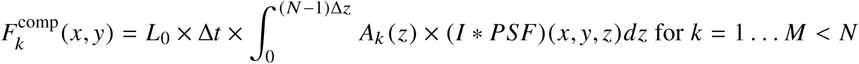

where *A_k_* (*z*) is a function that describes how the image intensity profile along the *z*-axis is combined into the single frame *k*. Note that this scheme is compressed in the sense that it requires *M* < *N* frames. This expression can be approximated and discretized as:

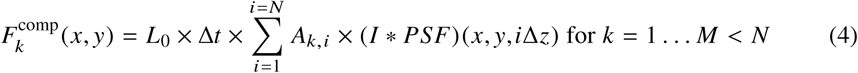

In the equation above, the matrix **A** is the discrete counterpart of *A_k_* (*z*) (*k* = 1 … *M*) and describes how the image intensity from each of the *N z*-planes of the stack is combined into a single value. Note that in the case where the measurement matrix is the identity matrix (*A_k,i_* = *δ*(*i*,*k*) with *N* = *M* and *δ* the Kronecker symbol), this scheme reduces to the plane-by-plane imaging scheme of equation (3) (Figure 1.b).

The connection with the compressed sensing setting outlined in Section 2.1 is immediate. For any fixed (*x, y*) location, 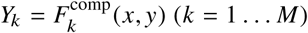 defines a *M* × 1 vector *Y*, that is a set of compressed measurements obeying *Y* = **A***X*, where *X* = *Ĩ* is the *N* × 1 vector corresponding to the intensity profile along the *z* axis convolved with the PSF and sampled every Δ*z*, i.e.:*Ĩ_i_* = (*I* * *PSF*)(*x, y*,*i*Δ*z*) for *i* = 1 … *N*. Thus, the matrix formulation of compressed sensing for a given (*x, y*) location is: *F*^comp^ = **A***Ĩ*. Applying the results mentioned in Section 2.1, under suitable conditions, approximate recovery of *Ĩ* should therefore be possible from the compressed measurements *F*^comp^.

### 2.3. Physical implementation

In practice, our microscopy system achieves axial compression in the following way. During each camera exposure, the stage is swept at constant speed across the entire z-range of the volume to be imaged (Figure 1.c):

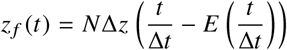

In this way, each pixel (*x, y*) of the camera records an integration of the emitted fluorescence along the *z* axis. During this axial sweep, the excitation laser intensity at the sample, *L*(*t*) = *L*_0_*T*(*t*)is modulated over time, such that:

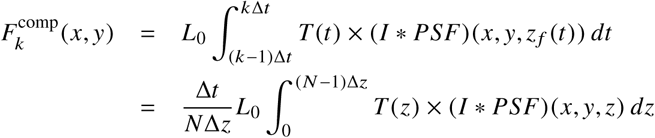

is an average of the fluorescence distribution along *z* weighted by the modulated excitation light intensity. In the discrete approximation, we have:

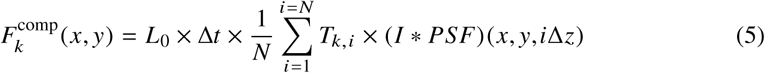

where *L*_0_ × *T_k,i_* is the laser power applied during frame *k* when the focal plane is at *z* = *i*Δ*z*. This modulation obeys a user-defined pattern specified by the *k*-th row of the measurement matrix **A**, as in equation (4) (Figure 1.c), i.e. we set **T** = *N***A**.

This procedure is repeated for all *M* rows of the measurement matrix, resulting in *M* compressed 2D images 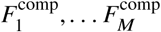. For a given constant exposure time Δ*t*, the acquisition speedup compared to plane-by-plane imaging is thus simply given by the ratio: *κ* = *N/M*. Figure 1.c jointly shows the *z* position of the stage and the illumination intensity for a measurement matrix **A** consisting of a truncated Fourier basis (shown in Figure 1.d).

A major advantage of this scheme is that it can be implemented without modification of the light path and simply requires a synchronization of two microscope components: (1) the *z*-piezo, which allows precise control of the axial focus (sample or objective) and (2) the AOTF (acousto-optic tunable filter) which allows to precisely and rapidly modulate the excitation light intensity transmitted to the sample (**T**). Instead of using an AOTF, it is in principle possible to modulate the intensity of the light source directly.

Note, that since the optical coding is performed by light modulation, this setup leaves absolute freedom of choice for the measurement matrix **A**, as long as its values are all positive. For example, random sensing matrices, which allow compressed sensing reconstruction for images sparse in any transform basis Ψ (universality property), can be implemented in a straightforward manner. In this paper, we choose a Fourier matrix, which is an optimal sensing matrix for images that are sparse in the direct spatial domain, taking the *M* first rows of the matrix, from low to high frequencies (Figure 1.d). Importantly, we linearly scale the matrix **A** such that all its values fall between 0 and 1/*N*. This ensures that during each frame *k*, the sample receives a light dose of 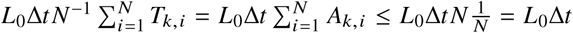, i.e. less or equal to the dose received in plane-by-plane imaging. Therefore, the total light dose received by the sample during a compressed imaging acquisition with *M* frames is at least *κ* = *N/M* times less than in the plane-by-plane acquisition, where *κ* is the compression ratio with respect to the plane-by-plane, Nyquist sampling.

### 2.4. Sparsity prior and PSF model

In this paper, we assume for simplicity that the 3D distribution of fluorescent structures is sparse in the spatial domain, but we take into account the 3D blurring caused by diffraction. This is done by incorporating a model of the 3D PSF into the *W* × *N* redundant dictionary Ψ, such that the 3D image can be modeled as: *X* = Ψ*α*, where *α* represents the 3D distribution of fluorescent structures and is assumed to be (approximately) sparse. In practice, we first measure the empirical PSF of the microscope using a conventional plane-by-plane *z*-stack and derive one cropped 2D image in the (*x, z*) plane. We then build the dictionary Ψ: each element of the dictionary is defined as a translation in the (*x, z*) plane of the empirical PSF within a given 2D reconstruction window. Thus, the dictionary is a collection of PSFs at various locations. Before reconstruction, both the compressed stacked and the elements of the dictionary are flattened into 1D vectors. Therefore, although our scheme performs compression only along the z axis, the reconstruction algorithm incorporates a 2D sparsity prior [26].

### 2.5. Numerical implementation

The reconstructions are performed using a custom port in Python of the previously published `SPIRAL–TAP` algorithm [24], which solves the optimization problems (1) or (2) above. Our port is publicly available, together with sample data and analysis scripts (see Section Software and data availability).

In practice, performing a reconstruction on a full 3D (or even 2D) image is not computationally tractable. Since we use a PSF model with a restricted spatial support, we assume that two (*x, y*) positions located farther apart than the characteristic width of the PSF are independent and reconstruct them in parallel on a computing cluster. We then calculate a single 2D image by averaging small overlapping chunks, and stack them together to obtain a 3D image.

## 3. Compressed imaging on simulated images

### 3.1. Generation of test images and metrics

We first tested our compressed sensing approach on simulated images. For this purpose, we generated a series of 100 synthetic images, termed ground truth (**GT**). The images contained features at different scales and sparsity levels (Figure 2a, top) along the compression axis (*z* axis). The images are scaled so that the pixel of highest intensity had a value of *I_max_* = 10,000 counts. For simplicity, the effects of diffraction blurring is ignored here.

**Fig. 2.**
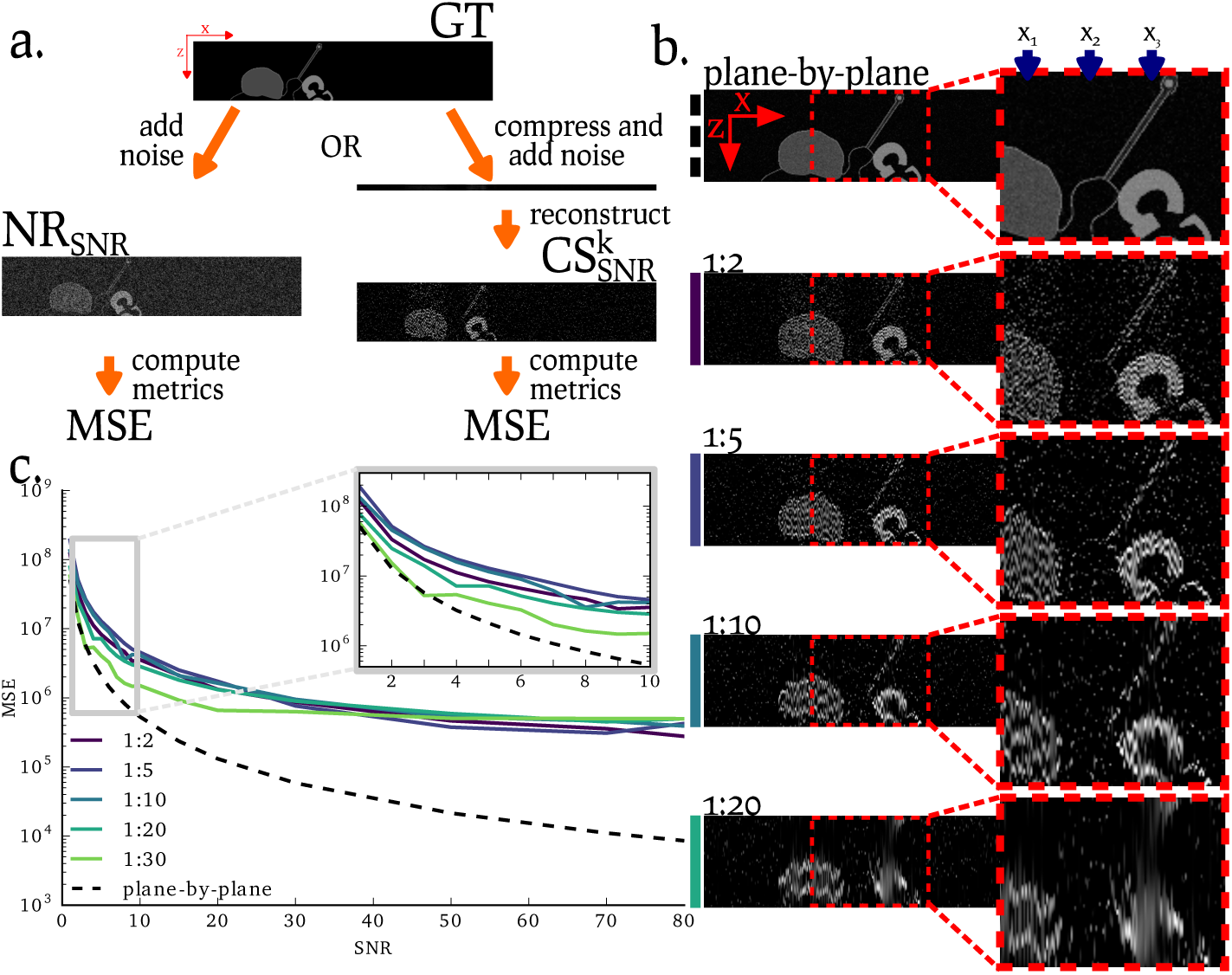
Simulations comparing the compressed sensing scheme with the traditional plane-by-plane scheme under various compression ratios and SNR. **(a).** Principle of the simulation: from a generated ground truth image (**GT**) and a specified SNR, (left) a noisy reference (**NR_SNR_**) is generated by adding Poisson and Gaussian noise to the ground truth. In parallel (right), the ground truth is compressed (along the *z* axis) with a compression ratio *κ* and an equivalent amount of noise is added at the same time as the compression (see main text for details). The compressed images are further decompressed (image 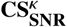) and the mean square error (MSE) is computed with respect to the ground truth. **(b.)** Examples of simulated images in the (*x, z*) plane. From top to bottom: (plane-by-plane) **GT**, (1:2) to (1:20) 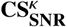 image reconstructed at a SNR of 20 and with a compression ratio of 1:2 to 1:20. The blue arrows represent lines of low sparsity, high sparsity and medium sparsity (respectively *x*_1_, *x*_2_ and *x*_3_). (**c.**) quality of the reconstruction (assessed by the MSE with respect to **GT**) for various SNR and compression-ratios. The dotted line is the MSE of the noisy reference **NR_SNR_** with respect to the ground truth **GT**. Inset: close-up on the low SNR region (SNR=1-10).

Next, to simulate an acquisition in realistic conditions in the conventional plane-by-plane scheme, we corrupted the ground truth images **GT** using Poisson noise (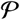) and additive half-normal noise 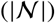. This resulted in noisy reference images (**NR_SNR_**) for different SNRs (see below): 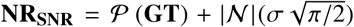 where *σ* corresponds to the expected number of background photons to achieve a given SNR, that is: *σ* = *I_max_/SNR*.

In parallel, we computed compressed versions of the same ground truth image by applying the measurement matrix **A** to **GT** (for different compression ratios *κ*, i.e. varying numbers of rows of **A**), and subsequently applied Poisson and additive Gaussian noise as for the images in the plane-by-plane acquisition 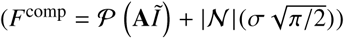.

Then, we computed reconstructions from these noisy compressed images, and denote the resulting 3D images as 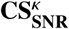. The reconstructions were performed without a PSF model, thus assuming sparsity of the reconstructed image and using Ψ = **I_N_**. The SNR was computed as the mean of the non-zero pixels of the ground truth image divided by the mean of the additive noise (Figure 2.b).

Finally, we quantified reconstruction quality by computing the mean square error 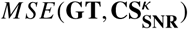 between the ground truth images **GT** and the compressed sensing reconstructions 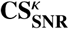. For comparison, we also computed the *MSE* between the ground truth images and the plane-by-plane acquisition image obtained for the same SNR, *MSE*(**GT, NR_SNR_**).

### 3.2. Results on simulated images

We first compare reconstructions for a fixed SNR=20 and increasing compression ratios *κ* (Figure 2.b). At low compression ratios (1:2, corresponding to a two fold speedup compared to the plane-by-plane acquisition), all features of the simulated images are accurately reconstructed, including both high and low frequency details. Minor artifacts are visible in the regions of moderate sparsity (arrow *x*_1_). As the compression ratio increases, fine features are progressively lost (arrow *x*_2_), whereas larger objects remain visible at their correct location (arrows *x*_1_ and *x*_3_). At high compression ratios, the reconstructed intensity significantly diverges from the ground truth. Nevertheless, this first example illustrates that object positions and shapes can be approximately reconstructed from compressed images with high compression ratios, even when the ground truth images exhibit quite variable levels of sparsity.

We then evaluate the influence of different noise levels on reconstruction quality by computing the MSE for SNR ranging from 1 to 80 and for compression ratios ranging from 1:2 to 1:30 (Figure 2c). At high SNR, the noisy plane-by-plane reference (**NR_SNR_**) exhibits a much lower MSE than the compressed sensing reconstruction. However, as mentioned in the introduction, compressed sensing is most useful for conditions in which photodamage and/or acquisition speed are limiting, i.e. for low SNR images. Figure 2c shows that for low SNR (≤ 15), the MSE of the noisy reference and the reconstruction become close. This suggests that our compressed sensing approach can recover images of similar quality as plane-by-plane imaging, but with a considerable reduction of acquisition time and light exposure. Equivalently, using the same acquisition time and light exposure as in plane-by-plane imaging, compressed imaging enables reconstructions of higher quality images as measured by the MSE metric.

Thus, our simulation results suggest that axially compressed sensing acquisition might be a worthwhile alternative to plane-by-plane imaging for faster and less phototoxic 3D microscopy.

## 4. Compressed lattice light sheet imaging

### 4.1. Lattice light sheet implementation

Having demonstrated our technique on simulated images, we implemented our compressed sensing scheme on a lattice light sheet microscope (LLSM). In a light sheet microscope [27], one objective is used to produce a very thin sheet of light that illuminates the sample at a 90° angle with the axis of another objective used for detection. Due to its excellent axial resolution, a lattice light sheet microscope is an ideal candidate to implement our compressed sensing scheme.

In LLSM, the focus is adjusted by moving the light sheet (using a scanning galvanometer mirror) and the focus of the observation objective (which is mounted on a piezo stage) in a synchronized manner across the sample (Figure 3a). For the compressed sensing acquisitions, we modified the software generating the FPGA control command (Coleman Technologies) in order to synchronize the motion of the sheet and the observation objective with a custom light modulation during a single camera exposure. Our modified software also allows to load a predefined measurement matrix **A** and to set exposure parameters.

**Fig. 3.**
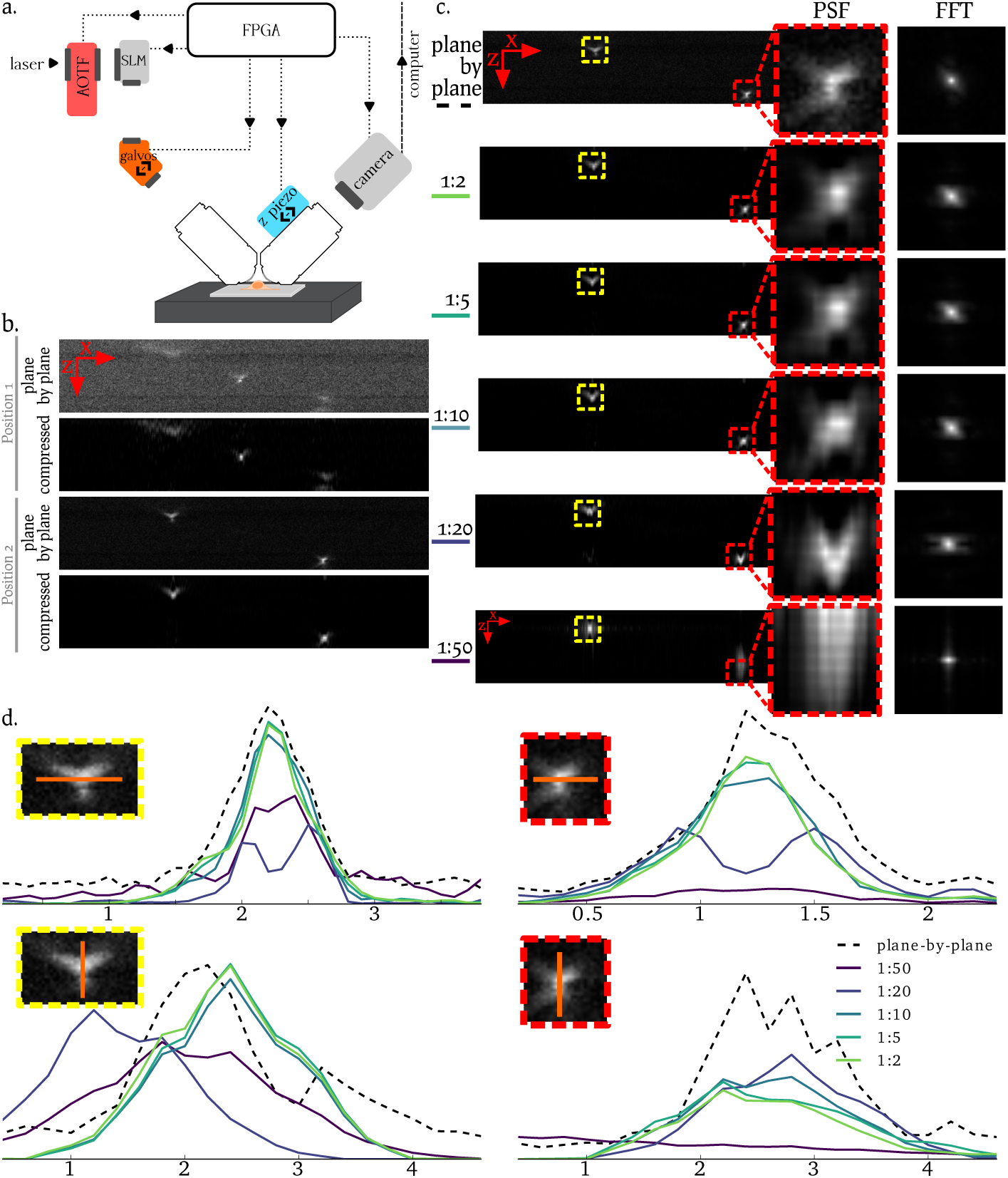
Compressed imaging reconstructions of fluorescent beads acquired with a lattice light sheet microscope. **(a).** Principle of a lattice light sheet microscope: two objectives at a 90°angle are used to observe the sample. The light sheet is generated through a spatial light modulator (SLM) and associated optics. The focus is adjusted by a coordinated move of the *z* piezo (that translates the observation objective) and of the *z* galvo (that translates the light sheet). Synchronization is achieved by a FPGA (Field-Programmable Gate Array). **(b).** Sample reconstructions (*compressed*) in the (*x, z*) plane from a 1:10 compressed acquisition and the corresponding acquisition in the plane-by-plane imaging scheme for two *y* positions (termed position 1 and position 2). **(c).** Compressed imaging sample reconstructions at increasing compression ratios (from 1:2 to 1:50) compared to the plane-by-plane scheme (top). For each reconstruction, a close-up on the PSF is displayed (*PSF* column) and the 2D frequency content of the PSF is displayed next to the PSF (*FFT* column) **(d).** Line profile across the two highlighted PSF (in yellow and red, respectively left and right). *top* Close-up along the *x* axis, *bottom* close-up along the *z* axis. The colors correspond to different compression ratios and the dotted line to the plane-by-plane reference. Horizontal axis in *μ*m. A field of view in the (*x, z*) plane is 50x20 *μm* (512x101 px). Full 3D reconstructions are shown in Supplementary Movies.

For the experiments reported below, we performed two sets of acquisitions: (i) one plane-by-plane acquisition for reference, and (ii) one compressed sensing acquisition. The plane-by-plane acquisition was performed by setting the measurement matrix **A** equal to the identity matrix in the acquisition software (**A** = **I_N_**) (Figure 1.b), in order to facilitate comparisons with the compressed imaging scheme and acquiring a z-stack with 101 frames at a fixed exposure time of Δ*t*=100 ms or 200 ms depending on the sample (see below). For the compressed sensing acquisition, we used a Fourier measurement matrix (Figure 1.d) with the appropriate scaling (see Section 2.3) to ensure that the light dose delivered to the sample for each camera exposure was equal to or lower than in the plane-by-plane acquisition. We acquired 50 frames 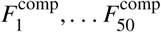 with the same exposure time Δ*t*, i.e. corresponding to a compression ratio of 1:2. To analyze higher compression ratios, we simply considered subsets of these frames 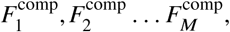 with *M* < 50. The highest compression ratio, 1:50, was obtained by keeping only the two first compressed frames 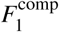 amd 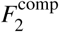. Varying the compression ratio in this manner allowed us to assess the minimum number of measurements required to obtain an acceptable reconstruction quality.

### 4.2. Results on fluorescent beads

We first imaged fluorescent beads (Tetraspeck 100 nm beads) on a glass coverslip. The exposure time for a single frame was set to Δ*t* = 100 ms and the stage was scanned over a 20 *μm* range (i.e. Δ*z* = 0.2*μm*).

Figure 3.c compares a 3D image of two fluorescent beads obtained through plane-by-plane imaging to 3D images reconstructed from the compressed acquisition for increasing compression ratios *κ* (from 1:2 to 1:50). It is apparent that qualitatively, the bead signal is reconstructed in the correct position for all compression ratios, including the highest (1:50), which corresponds to only 2 frames (instead of 101 in the plane-by-plane acquisition). As the compression ratio is increased from 1:2 to 1:50, the reconstructed images of the two beads (i.e. PSFs) progressively deteriorate and became significantly distorted for compression ratios ≥ 1:20. However, up to a compression ratio of 1:10, all the beads and the fine features of the PSF shape are properly and accurately reproduced (Figure 3.b). This is also apparent in the line profiles in Figure 3.d. Thus, this experiment illustrates that high quality reconstruction of 3D images is possible in compressed imaging using 1s of total acquisition time, compared to 10s in the plane-by-plane imaging scheme, thereby representing a 10 fold speedup.

### 4.3. Results on fixed cells

Encouraged by these results on beads, we proceeded to imaging fixed cells. Mouse embryonic stem cells (mESCs) were seeded on glass coverslips and fixed in 4% paraformaldehyde (PFA). The cells were then stained for actin with a fluorescently labeled probe (phalloidin-RFP) and imaged in an oxygen-scavenging medium. The camera exposure time was set to Δ*t* = 200 ms and the *z* scanning range to 20 *μm* (Δ*z* = 0.2*μm*).

Results are shown in Figure 4.a, where the 3D reconstruction of a sample imaged with 1:5 compression (right) is compared to the plane-by-plane acquisition (left). It is apparent that in both lateral (*x, z*) and axial (*x, y*) sections, fine and large details are successfully reconstructed. Although some high frequency details are lost in the (*x, z*) plane, most features are faithfully reproduced. This observation is further confirmed by the maximum intensity projection of the reconstructed stacks at various compression ratios (Figure 4.b). For compression ratios up to 1:5, the reconstructions show intensity profiles very similar to those of the plane-by-plane reference image. At higher compression ratios, however, significant artifacts become visible along the *z* axis (profiles in Figure 4.a,b bottom). Artifact-free reconstructions with higher compression ratios might be possible using a number of possible improvements (see Discussion). Nevertheless, these results already indicate that our compressed sensing approach is a viable method to achieve substantial reductions of acquisition time (and light exposure) for 3D imaging of biological samples.

**Fig. 4.**
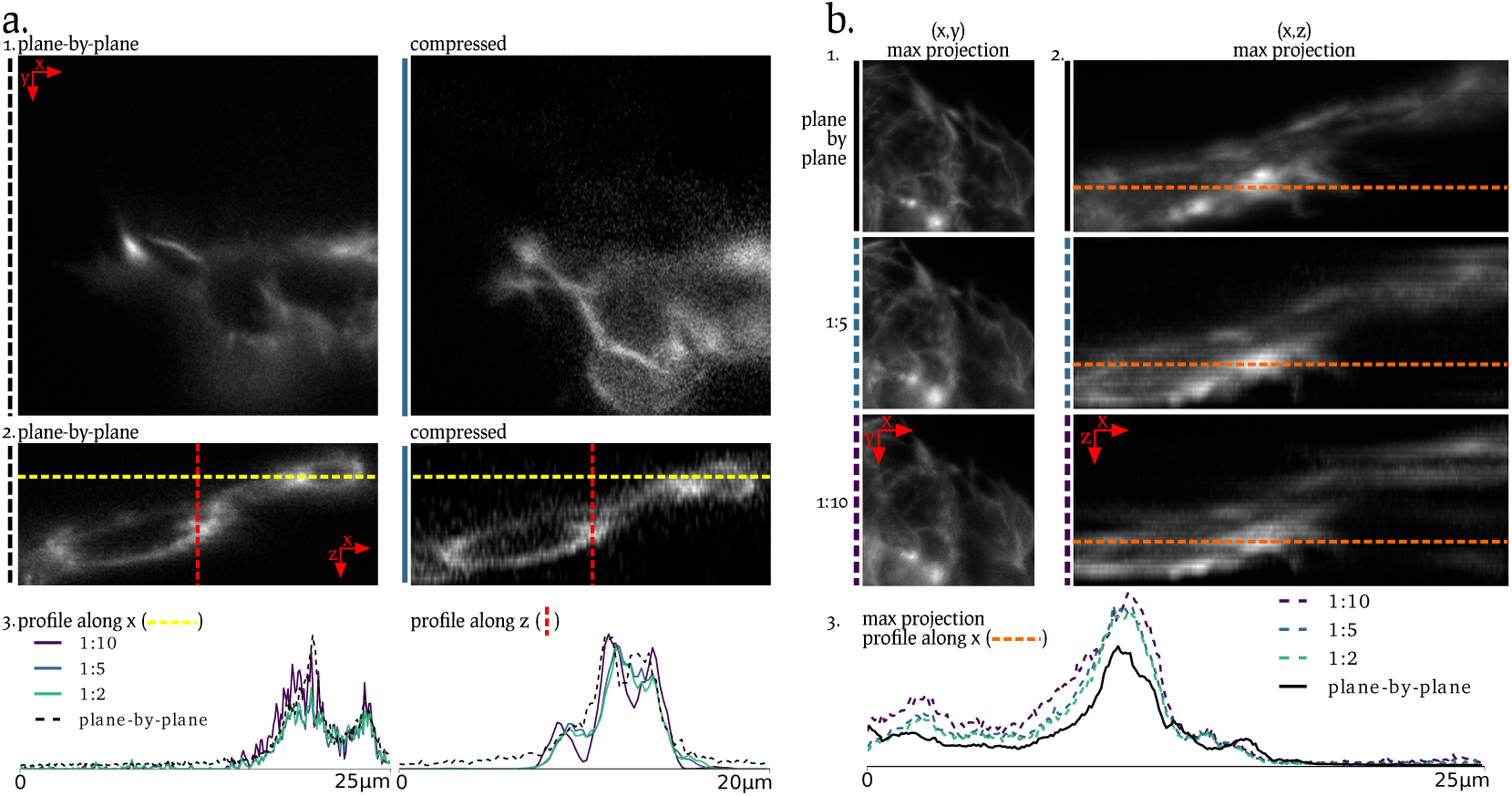
Compressed imaging reconstruction of phalloidin-RFP-stained, fixed mouse embryonic stem cells (mESCs) acquired with a lattice light sheet microscope. **(a).** Example reconstruction from a 5 fold compression ratio (right) and the corresponding plane-by-plane reference acquisition (left) (*1., top* in the (*x, y*) plane and (*2., middle*) in the (*x, z*) plane. Dotted lines indicate the location of the line profiles presented in (panel *3., bottom*) (left) profile along the *x* axis (yellow dotted line of panel 2.) for increasing compression ratios. (right) profile along the *z* axis (red dotted line of panel 3.) the curves are the average over 3 planes in the *y* dimension. **(b.)** Maximum intensity projection in the (*x, y*) plane (*1., left*) and in the (*x, z*) plane (*2., right*) of the plane-by-plane stack (top) and reconstructed stack at increasing compression ratios (two bottom pictures). The orange dotted line shows the location of the line profile displayed in panel *3., bottom*: line profile of the reconstruction at increasing compression ratios (dotted line) compare to the plane-by-plane reference (continuous line). A field of view in the (*x, z*) plane is 25x20 *μm* (256x101 px) and a field of view in the (*x, y*) plane is 25x25 *μm* (256x256 px).

## 5. Compressed epifluorescence microscopy

### 5.1. Implementation

We also implemented our compressed sensing scheme on a standard epifluorescence microscope, an almost ubiquitous instrument in cell biology labs. Since our imaging strategy relies only on the synchronization of the stage position and the light modulation, it is very versatile and suitable for a wide range of microscopes.

An epifluorescence microscope (Nikon Eclipse TI) equipped with an AOTF (AOTFnC-400-650-TN, AA Optoelectronics) and a *z* piezo stage (Nano-ZL 500, Mad City Labs) is controlled using an Arduino microcontroller (Genuino Uno) to synchronize the AOTF and the stage through their analog input, based on the camera fire signal (Figure 5.a). The microscope was controlled using MicroManager [28], and custom firmware was written for the Arduino, allowing for a software switch between plane-by-plane and compressed sensing acquisition. In practice, the measurement matrix **A** is first loaded to the Arduino, then the focus is adjusted in the plane-by-plane imaging mode and an acquisition sequence is set using a custom MicroManager plugin. Finally, the Arduino is switched to the compressed sensing mode and the images are acquired and handled using the usual MicroManager logic.

**Fig. 5.**
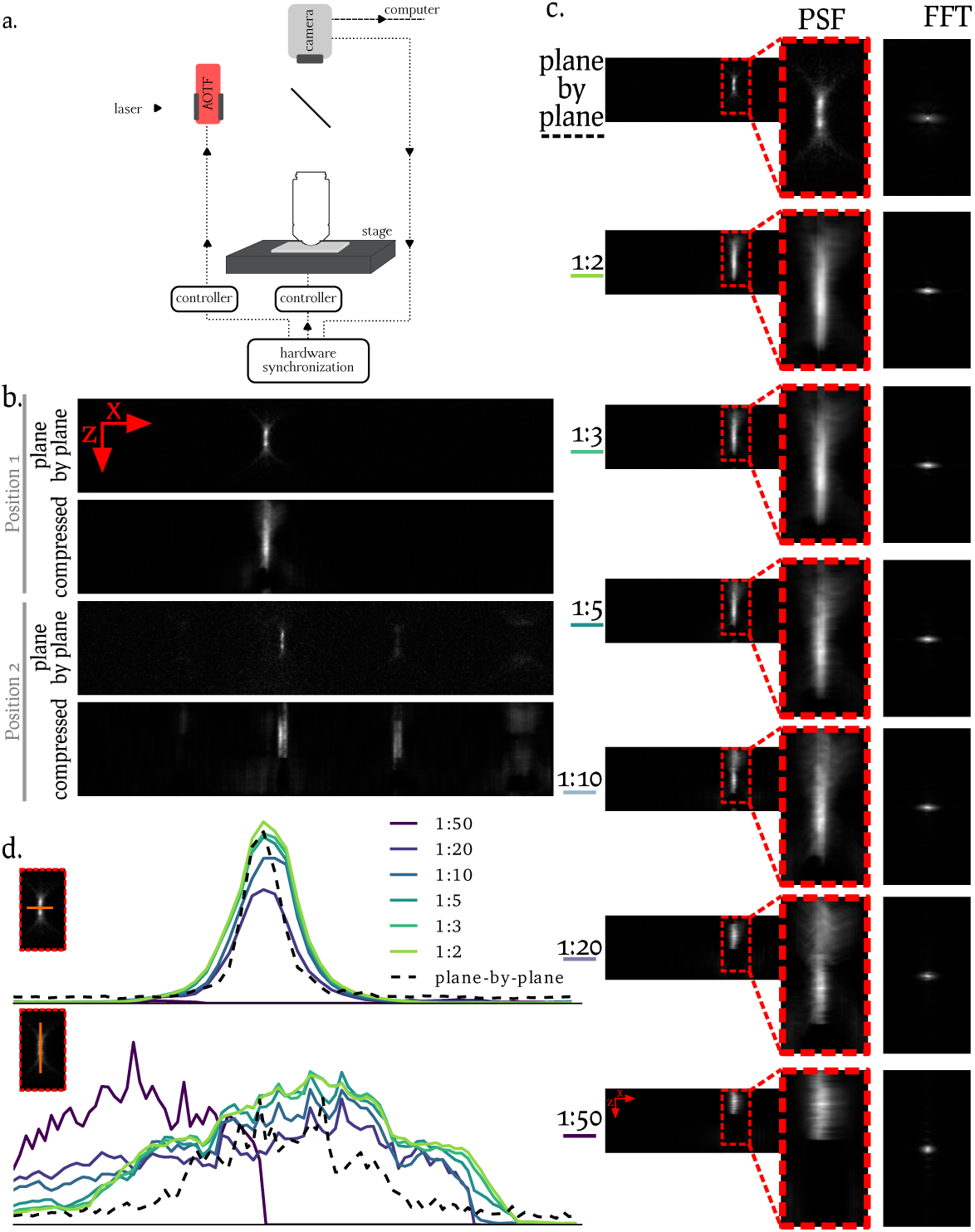
Compressed imaging reconstructions of fluorescent beads acquired with an epifluores-cence microscope. **(a).** Principle of the epifluorescence microscope: both the AOTF and the motorized stage in *z* are synchronized by hardware (Arduino). **(b).** Example reconstructions (*compressed*) in the (*x, z*) plane from a 1:10 compressed acquisition and the corresponding acquisition in the traditional imaging scheme (*plane-by-plane*) for two y positions (termed position 1 and position 2). **(c).** Sample reconstructions at increasing compression ratios (from 1:2 to 1:50) compared to the plane-by-plane imaging scheme (top). **(d).** *x* and *z* profiles of one selected PSF (highlighted in panel c) reconstructed from various compression ratios. The black dotted line represents the plane-by-plane reference. Full 3D reconstructions are provided as supplementary materials.

### 5.2. Results on fluorescent beads

We imaged fluorescent beads on a glass coverslip with an axial range of ∼ 100 *μ*m and an exposure time of Δ*t* = 200 ms. Results are shown in Figure 5.b-d for varying compression ratios *κ*. It is apparent from Figure 5.c that the reconstructed images qualitatively recover the beads position accurately for compression ratios up to 1:10. Interestingly, the reconstructed images display a decreased level of background noise, revealing the PSF shape of out-of-plane beads (i.e. beads in another *y* = *constant* plane) that are not visible in the noisy, plane-by-plane imaging reference (Figure 4.b). Although the location of the PSF in the reconstructed image is consistent with the plane-by-plane reference (Figure 4.d), the PSF in the latter is significantly sharper than in the reconstructed image. This might be due either to an insufficient synchronization of the AOTF with the stage or to an inaccurate PSF model used for the reconstruction. For higher compression ratios (≥ 1:20), the PSF is no longer well localized in *z*.

These results indicate that our compressed sensing approach can be successfully applied to an epifluorescence setup and achieve roughly ten-fold compression ratios. Given the large availability of epifluorescence microscopes, this shows that the benefits of our compressed imaging approach can be made widely accessible with little effort.

## 6. Discussion

In this paper, we present a new compressed sensing scheme for 3D fluorescence microscopy that can be applied to a wide range of microscopes. We validate our method through simulations, and demonstrate its feasability on a lattice light sheet microscope and on an epifluorescence microscope. We achieve reductions in image acquisition time and light exposure (compression ratios) of up to ten fold on both setups.

Most previously proposed compressed sensing strategies for biological microscopy require non-trivial modifications to microscope hardware, such as adding a digital micromirror device to the light path [13], or conjugating the camera with the back pupil plane to perform Fourier space imaging [12]. By contrast, our compressed sensing scheme is adaptable to any type of epifluorescence microscope, as long as the user can control both the stage position and the illumination intensity. Our lattice light sheet microscope implementation required software adjustments due to its complex nature, while implementation on an epifluorescence microscope required only one extra microcontroller (to ensure proper synchronization of camera, AOTF and stage). We provide full schematics of the Arduino setup, together with a MicroManager plugin, which allows to quickly switch between plane-by-plane imaging and the compressed sensing imaging scheme.

Since our approach modulates the light intensity before it reaches the sample it results in a reduced excitation light dose to the sample. This is of crucial importance, since fluorescence imaging causes damage to the sample, ranging from photobleaching of the fluorescent probes to various metabolic and developmental defects. Furthermore, the light dose delivered to the sample is inversely proportional to the compression ratio: a ten fold compression ratio yields a ten fold reduction of the light dose at the sample compared to plane-by-plane imaging. Since phototoxicity is a nonlinear effect [29], we expect our compressed imaging method to allow dramatic reductions in photodamage.

Another major benefit of our compressed imaging scheme is an increase in the temporal resolution of 3D imaging, thus allowing faster acquisition at a given SNR. For example, if plane-by-plane imaging requires 200 planes with 10 ms exposure each, i.e. a total of 2 s to acquire a given 3D volume, a compressed imaging scheme with a compression ratio of 10 will require only 200 ms for the same volume. This reduction in acquisition time should enable an equivalent increase in the temporal resolution of dynamic 3D microscopy of living biological samples. We anticipate that future work will build on the proposed approach to explore the potential of axially compressed imaging for faster 3D live cell imaging.

In this context, several challenges and perspectives for improvement are worth mentioning. First, better piezo hardware and/or calibration could reduce the mismatch between the theoretical sensing matrix and its experimental counterpart. Second, as in plane-by-plane imaging, our current method assumes that the imaged structure remains immobile throughout compressed acquisition of the 3D volume, which is rarely true in living samples. While sample movements can result in a blurred image or duplicated objects with traditional plane-by-plane imaging, it remains to be explored how these movements may distort reconstructed images in our compressed sensing scheme. As previously shown in the MRI field, methods to address these problems can be developed [26]. Third, computational strategies to accelerate image processing will be important to efficiently analyze the thousands of images generated in dynamic 3D imaging [30,31]. Finally, our method currently assumes sparsity in the image domain along the optical axis only and used a Fourier sensing matrix. Ten fold compression ratios were achieved on images with moderate degrees of sparsity, for which this sensing matrix is not optimal. We therefore expect that larger compression ratios can be achieved using 3D sparsity models better adapted to the imaged structures, e.g. using dictionary learning, or optimized sensing matrices [32, 34].

## 7. Conclusion

In this work, we demonstrate a widely applicable 3D compressed sensing scheme in which images are compressed along the *z* axis during acquisition. This scheme can be implemented with little or no hardware modification on a wide range of microscopes. We first validated the feasability of our approach under noisy conditions using simulations, and then demonstrated the method experimentally on a lattice light sheet microscope and on an epifluorescence microscope, where we achieved roughly ten fold imaging speedup. This approach can be used to increase the temporal resolution or extend imaging time (through a reduction in light dose) in 3D fluorescence imaging applications.

## Software and data availability

A Python port of SPIRALTAP [24] is available on GitHub https://github.com/imodpasteur/pySPIRALTAP (doi:10.5281/zenodo.439691. The datasets used in this work are available on the Zenodo repository (doi:10.5281/zenodo.439689) and the analysis scripts were deposited on GitHub https://github.com/imodpasteur/CompressedSensingMicroscopy3D (doi:10.5281/zenodo.439690). The MicroManager plugin and Arduino device adapter are available on https://github.com/imodpasteur/ArduinoCompressedSensing.

## Acknowledgments

We are very grateful to Zach Harmany who made available the SPIRALTAP software under an open source license and to Moran Mordechay for useful discussions on further optimization of the imaging scheme. We would like to thank the whole IMOD lab and Darzacq labs for insightful discussions and suggestions to this work.

This work used the computational and storage services (TARS cluster) provided by the IT department at Institut Pasteur, Paris.

